# The base-editing enzyme APOBEC3A catalyzes cytosine deamination in RNA with low proficiency and high selectivity

**DOI:** 10.1101/2021.11.26.470160

**Authors:** Aleksia Barka, Kiara N. Berríos, Peter Bailer, Emily K. Schutsky, Tong Wang, Rahul M. Kohli

**Author notes:** Co-first authors (Aleksia Barka, Kiara N. Berríos).

## Abstract

Human APOBEC3A (A3A) is a nucleic acid-modifying enzyme that belongs to the cytidine deaminase family. Canonically, A3A catalyzes the deamination of cytosine into uracil in single-stranded DNA, an activity that makes A3A both a critical antiviral defense factor and a useful tool for targeted genome editing. However, off-target mutagenesis by A3A has been readily detected in both cellular DNA and RNA, which has been shown to promote oncogenesis. Given the importance of substrate discrimination for the physiological, pathological, and biotechnological activities of A3A, here we explore the mechanistic basis for its preferential targeting of DNA over RNA. Using a chimeric substrate containing a target ribocytidine within an otherwise DNA backbone, we demonstrate that a single hydroxyl at the sugar of the target base acts as a major selectivity determinant for deamination. To assess the contribution of bases neighboring the target cytosine, we show that overall RNA deamination is greatly reduced relative to that of DNA, but can be observed when ideal features are present, such as preferred sequence context and secondary structure. A strong dependence on idealized substrate features can also be observed with a mutant of A3A (eA3A, N57G) which has been employed for genome editing due to altered selectivity for DNA over RNA. Altogether, our work reveals a relationship between the overall decreased reactivity of A3A and increased substrate selectivity, and our results hold implications both for characterizing off-target mutagenesis and for engineering optimized DNA deaminases for base-editing technologies.

## INTRODUCTION

Purposeful enzymatic transformations to nucleic acids play critical roles in a host of biological processes, but also pose potential risks when mistargeted. The challenge of discerning between different nucleic acid substrates is particularly important for the AID/APOBEC family of enzymes, the majority of which canonically catalyze the hydrolytic deamination of cytosine into uracil within single-stranded DNA (ssDNA).^1^ AID functions to drive antibody maturation through the mutation of host immunoglobulin genes, while APOBEC3 family members are antiviral restriction factors mutating retroviral genomes through targeted deamination of replication intermediates.^1^ Although such targeted DNA deamination is beneficial for immunity, AID/APOBEC-catalyzed deamination is also a major source of mutation in numerous cancer types, highlighting the consequences of aberrant or misregulated mutagenesis (**Figure 1a**).^2–5^

**Figure 1.**
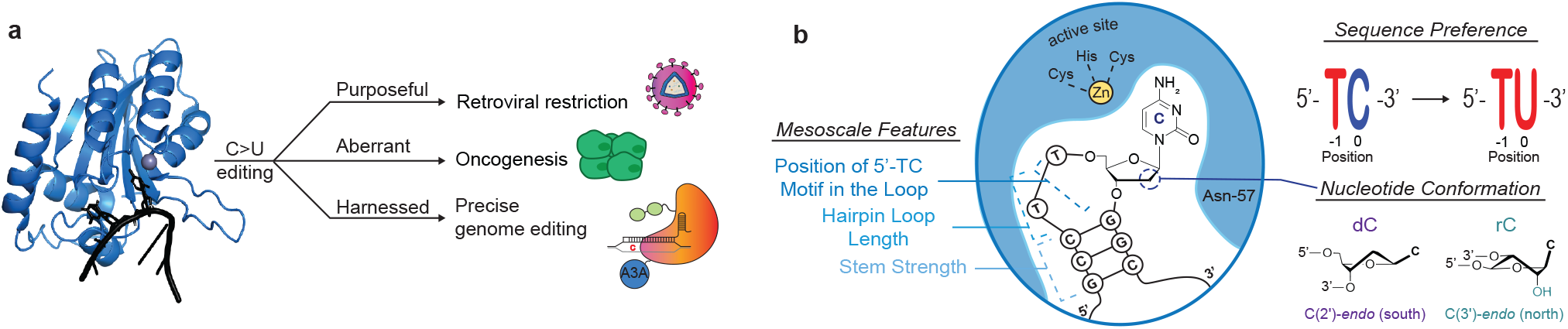
A3A mutagenesis and substrate selectivity features. (a) Schematic representation of the purposeful biological functions, pathological implications, and harnessed biotechnological applications of A3A-mediated deamination. Depicted on the left is the co-crystal structure (PDB: 5SWW) of A3A (blue) and ssDNA (black). (b) Schematic representation of the ssDNA-A3A complex with key features highlighted. The substrate is shown as a stem-loop corresponding to preferred substrate conformation. Conformation differences of the target nucleotide, sequence context preference, and mesoscale features are annotated.

Pro-oncogenic deamination can extend to either DNA or RNA. Pathological DNA deamination has been a particular area of focus for two human APOBEC3 family members – APOBEC3A (A3A) and its close relative APOBEC3B (A3B). Many cancer genomes harbor a clear signature of aberrant A3A or A3B activity, as evidenced by clusters of closely spaced and strand-coordinated cytosine mutations in TpC contexts, termed kataegis.^6–8^ In addition to shaping global mutational footprints, specific genes can be targeted. An intriguing example is offered by mutations in succinate dehydrogenase B (*SDHB*), which are frequently observed in leukemic T cells.^9^ Matched examination of the transcriptome and genome led to the discovery that A3A targets *SDHB* messenger RNA (mRNA) transcripts for deamination, and not the *SDHB* gene,^10^ highlighting how A3A activity on either DNA or RNA can have pathological consequences.

The question of substrate discrimination has taken on a new importance now that AID/APOBEC enzymes have also been harnessed for genome editing (**Figure 1a**). ‘Base editors’ utilize a catalytically-impaired Cas protein to direct a tethered DNA deaminase to a specific genomic locus to introduce a targeted single-base mutation.^11^ Base editors employing A3A are particularly appealing given their high efficiency; however, unwanted RNA off-target editing has also been observed.^12–15^ While an N57G mutation in engineered A3A (eA3A) has been shown to dampen RNA off-target activity while retaining on-target activity,^14^ the biochemical selectivity of eA3A on DNA versus RNA has not been directly explored.

Previous studies have shed some light into the potential mechanisms governing A3A’s substrate specificity. While there is speculation that RNA deamination may be a physiological role for A3A,^16^ the only APOBEC family member for which RNA activity is definitively established is APOBEC1.^17^ Nonetheless, most family members have been shown to bind RNA even tighter than DNA,^18^ adding to the enigma of nucleic acid selectivity. For several family members, a prominent role for the identity of the target cytidine nucleotide has been revealed. For AID, the substitution of a single ribocytidine (rC) in an otherwise DNA substrate reduces deamination at least 500-fold.^19^ Interestingly, APOBEC1 also shows a marked preference for DNA,^19^ a feature which may reflect its ancestral functions.^20^ While these studies offer important precedents, it remains unknown if the target cytidine nucleotide plays a similarly large role in governing nucleic acid selectivity by A3A.

Further complexity arises from genomic and structural studies that highlight unique aspects of A3A catalysis that distinguish it from other family members. In addition to the known TpC preference, secondary structure features have been found to significantly impact substrate selectivity for A3A (**Figure 1b**). The solution of a DNA-bound crystal structure of A3A revealed that ssDNA adopts a U-shaped conformation when bound in a catalytically competent orientation.^21^ This observation provides a rationale for results from genomic studies focused on analyzing mesoscale level (~30 base pair) features that are preferentially targeted by A3A in genomic DNA **(Figure 1b)**. These sequences are characterized by the following mesoscale features: the formation of stem-loop structures, the positioning of the cytosine base at the 3’ end of a 3-5 bp loop, and a strong stem.^22^ Biochemical and transcriptome-wide studies have also demonstrated similar stem-loop preferences with RNA;^23,24^ however, no study has yet to look at matched DNA and RNA substrates to assess the relative importance of these features in dictating selectivity.

In this report, we were motivated to decipher the mechanistic basis for nucleic acid discrimination by A3A given the enzyme’s role in immunity, cancer biology, and genome editing. We first characterize the role of the target cytidine in DNA/RNA selectivity, showing that the presence of a single 2’-hydroxyl group at the sugar of the target nucleotide markedly decreases reactivity. We then establish the role of neighboring bases and secondary structure as deterministic features of DNA versus RNA selectivity with both A3A and engineered eA3A. Overall, our results highlight a tradeoff between efficiency and selectivity, where DNA deamination is highly preferred and less selective while RNA deamination is less preferred but highly selective. Our findings provide new mechanistic insights into A3A mutagenesis and offer guidance for evaluating alternative A3A variants for biotechnological applications.

## RESULTS AND DISCUSSION

### Target nucleotide as a major selectivity determinant for deamination

RNA and DNA are distinguished by a single 2’-(*R*)-hydroxyl group (2’-OH) present on the ribosyl sugar of each nucleotide in RNA but not in DNA (**Figure 1b**). Enzymes that differentially act on DNA versus RNA have been shown to be influenced by the existence of the 2’-OH in different manners. For example, the base excision repair enzyme uracil DNA glycosylase (UDG) uses a steric exclusion mechanism to target DNA over RNA, and a similar ‘steric gate’ is employed by DNA polymerases.^25,26^ By contrast, a 2’-OH can alter favored nucleotide conformations which can influence selectivity independent of direct steric interactions. Differential sugar pucker dictated by the 2’-OH governs selectivity with RNA ligase, a mechanism that has also been shown to also apply to AID and APOBEC1.^19,27^

We posited that for A3A, in a manner akin to its AID/APOBEC relatives, the 2’-OH of the target cytidine itself might play a prominent role in substrate discrimination. In order to isolate this feature, we designed a 35 bp DNA substrate (S35-dC) and its associated product (S35-dU), along with matched chimeric versions that contained a single ribocytidine (rC) or ribouridine (S35-rC and S35-rU) within the otherwise DNA backbone. The base (^x^C) was embedded in an ATT<u>^x^C</u>AAAT sequence context, which includes the preferred TpC context for A3A and could newly introduce a cleavage site for the restriction enzyme SwaI upon successful deamination (**Figure 2a**). We first validated that upon duplexing, S35-rU could be cleaved as efficiently as S35-dU by SwaI, offering a facile means to track deamination. We next reacted the S35-dC or S35-rC substrate with serial dilutions of A3A, duplexed, and analyzed for product formation by quantification of the cleavage product (**Figure 2b**). Our *in vitro* assay revealed that the simple addition of a single 2’-OH reduced A3A efficiency by 110-fold (**Figure 2b-c**), indicating that the identity of the target nucleotide itself contributes substantially to substrate selectivity.

**Figure 2.**
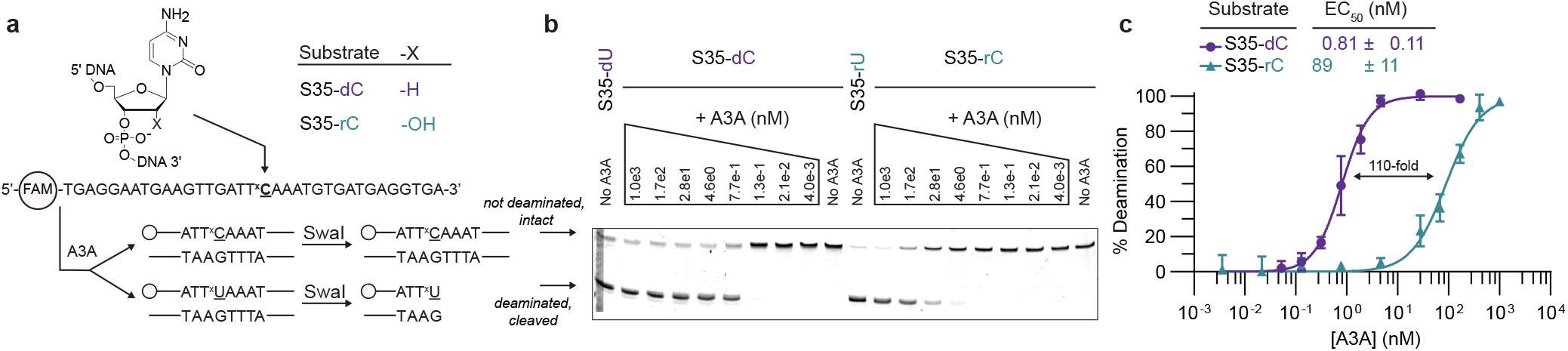
The addition of a single 2’-OH in the target cytosine decreases deamination efficiency. (a) Top – Chimeric substrate design. Target cytosine is either not modified (dC) or modified to have a 2’-OH substitution (rC) and embedded in an otherwise DNA backbone, resulting in S35-dC and S35-rC oligonucleotides, respectively. Bottom – Sequence of oligonucleotides and SwaI assay diagram. 5’-FAM-labelled S35-dC and S35-rC oligonucleotides have the target cytosine embedded in a TT<u>^x^C</u>AAA sequence context. After reaction with A3A, the oligonucleotides are duplexed to a complementary strand and digested with SwaI. SwaI cleaves the deaminated product but does not cleave the non-deaminated substrate. (b) A3A titration. Oligonucleotides (100 μM) are reacted with 6-fold dilutions of A3A (from 1 μM to 4 pM from left to right) for 30 min at 37 °C. Imaging of the fluorescent oligonucleotides in a representative denaturing polyacrylamide gel is shown. For each substrate, the leftmost lane includes a product control, and the rightmost lane includes a substrate control not treated with A3A. (c) Percent deamination is plotted for S35-dC (purple) and S35-rC (teal) as a function of A3A concentration and normalized to product controls. Data represent at least three independent replicates with mean and standard deviation plotted. Product formation was fit to determine the EC_50_, the enzyme concentration required to convert half of the substrate under the assay conditions.

The reduction in deamination efficiency upon the addition of a single hydroxyl to a DNA oligonucleotide is particularly notable given that the structure of DNA-bound A3A suggests that a 2’-OH could be accommodated without major steric conflicts (**Supplementary Figure 1**).^21^ This result suggests the possibility that the influence of the 2’-OH on sugar pucker, rather than on sterics, could alter the efficiency of deamination. Notably, the impact of the 2’-OH on A3A activity is closer to the level of discrimination observed with APOBEC1 (100-fold) rather than AID (500-fold), an observation consistent with the detection of higher levels of edited RNA via A3A, but not by AID.^19,28,29^

### Sequence context and secondary structure impact nucleic acid selectivity

While our initial results suggest that the role of the target base is a consistent feature of nucleic acid selectivity across the AID/APOBEC family, A3A is specifically distinguished from other family members by the prominent impact of mesoscale features in selective deamination of DNA.^22^ We therefore hypothesized that secondary structure and sequence context might similarly affect A3A’s selective deamination of DNA versus RNA.

To test this hypothesis, we moved from the chimeric substrates containing a single rC to more complex sequence-matched substrates composed entirely of DNA or RNA. To this end, we designed a long 720-mer substrate containing cytosines in diverse sequence contexts and potential secondary structure elements (**Supplementary Figure 2a-b**), including the SDHB hotspot which has been implicated in RNA editing.^10^ Importantly, the substrate was designed lacking Cs in the 5’-end and Gs in the 3’-end of the sequence to allow unbiased amplification of the sequence by either PCR or RT-PCR after reaction with A3A. The resulting amplicons could then be examined for deamination at specific sites by analyzing changes in restriction enzyme digestion contexts, or across the whole amplicon by next-generation sequencing (NGS) (**Figure 3a**). Specifically, detection of deamination at the SDHB target loop was possible using the restriction endonuclease ClaI. The intact (non-deaminated) site is in a 5’-AT<u>C</u>GAT-3’ context that can be cleaved by ClaI, while the deaminated product is not, offering a facile quantitative means to track deamination at this site (**Supplementary Figure 3a**). Using the matched 720-mer sequences, we performed deamination with serial 10-fold dilutions of A3A (6 μM to 6 pM) and analyzed for resistance to ClaI cleavage. For the ssDNA substrate, with 0.6 nM A3A, the majority of the substrate was deaminated at the SDHB site (**Figure 3b**). By contrast, comparable deamination was observed in the RNA substrate only at higher concentrations of A3A. We quantified deamination across replicates to determine the EC_50_, here defined as the amount of enzyme necessary to deaminate half the substrate at this site. This analysis determined that deamination at the SDHB site was 94-fold less efficient in RNA than in DNA (**Figure 3b**). Thus, even for this robust RNA deamination target, DNA deamination efficiency is substantially higher. We extended this analysis to a second restriction enzyme-responsive site within the amplicon, where consecutive deamination events at a 5’-T<u>CC</u>AAA site, converting it to 5’- T<u>TT</u>AAA, could be detected via digestion with DraI (**Supplementary Figure 3a**). The preference for DNA deamination was further evident at this site, as consecutive deamination in ssDNA was nearly complete with ≥6 nM A3A, while it was not detected on RNA even with maximal A3A (6 μM) (**Supplementary Figure 3b**).

**Figure 3.**
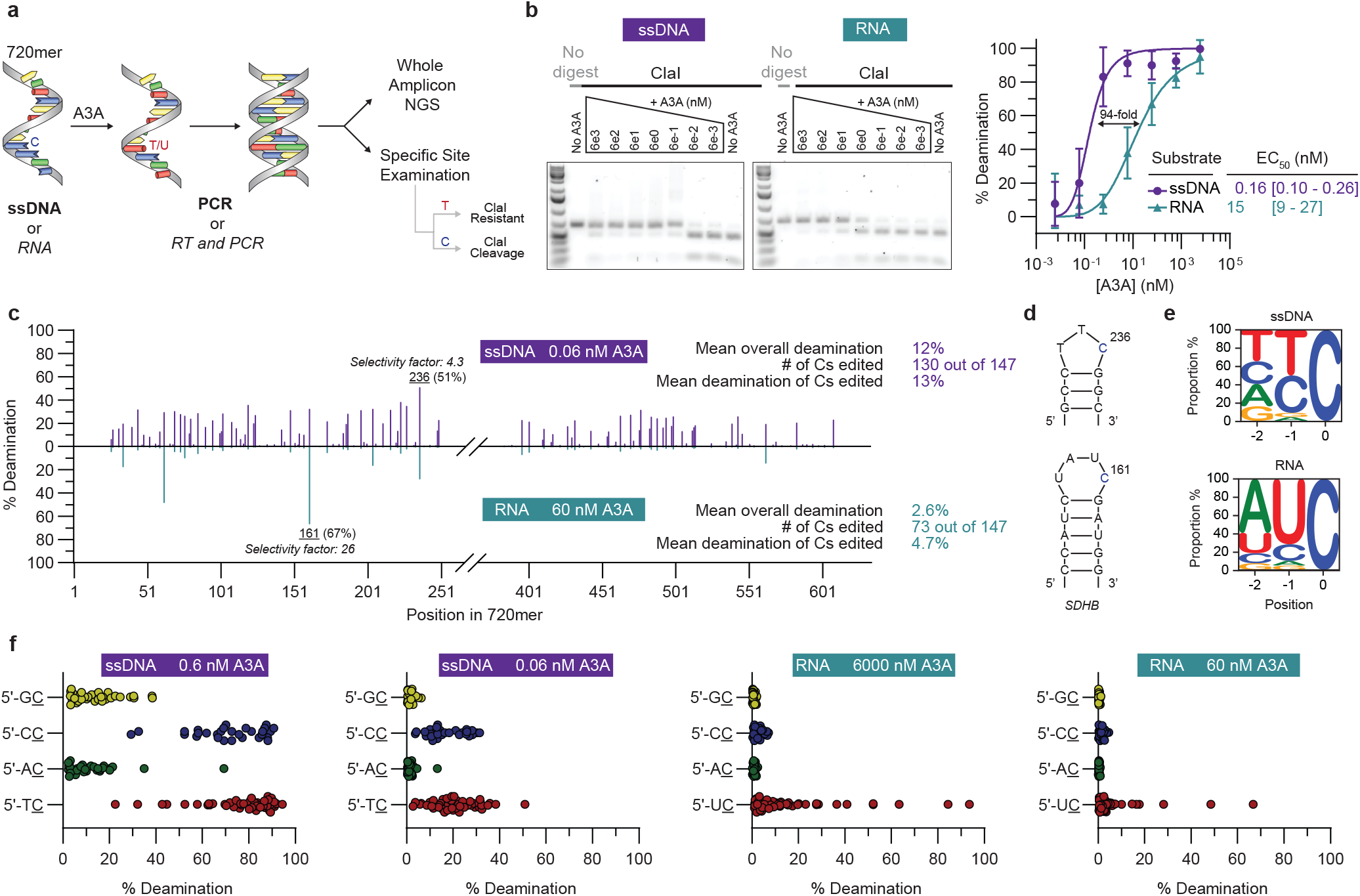
A3A activity on long, single-stranded substrates with matched sequences. (a) 720mer assay diagram. Sequence-matched ssDNA and RNA substrates were reacted with A3A. The samples were amplified by PCR (ssDNA) or RT-PCR (RNA). Amplified products were then subjected to site-specific examination via the use of restriction enzymes, such as ClaI, or to whole amplicon Next-Generation Sequencing (NGS). ClaI cleaves non-deaminated substrates but not the deaminated products. (b) Left – Representative gels of A3A titration. 10 ng of ssDNA and RNA substrates were reacted with 10-fold dilutions of A3A (6 μM to 6 pM, left to right) for 30 min at 37 °C. Following amplification, amplicons were digested with ClaI and imaged on a 1.5% agarose gel. Right – Quantification of percent deamination at the SDHB site for ssDNA (purple) and RNA (teal) as a function of A3A concentration. Data represent four independent replicates with mean and standard deviation plotted. Product formation was fit to determine the EC_50_. (c) Base resolution map showing percent deamination vs. the position of cytosines across the 720mer as per NGS analysis. ssDNA data from reaction with 0.06 nM A3A is shown in purple above the axis, while data from RNA reacted with 60 nM A3A is shown in teal below the axis. The middle 134 bp are not included in the analysis due to the limitations of paired-end sequencing. Data represent the mean deamination at each position from two independent experiments with results from individual amplicons provided in **Supplementary Table**. The most heavily deaminated cytosine for each substrate is labelled with its position in the 720mer, selectivity factor, and percent editing. (d) Schematic representation of the stem-loop structures of the most heavily deaminated cytosine for each substrate. Top – cytosine at position 236 in ssDNA. Bottom – SDHB site; cytosine at position 161 in RNA. (e) Sequence logos of editing sites for ssDNA and RNA samples after correcting for background editing levels. Position 0 represents the target C. (f) Jitter plots showing percent deamination for substrates reacted with different concentrations of A3A, separated by sequence context, highlighting the higher proficiency and lower specificity for deamination of ssDNA versus RNA.

To profile substrate discrimination more rigorously, we next quantified deamination via NGS, analyzing a total of 146 cytosines across the amplicon using 250 bp paired-end reads. Notably, our assay was robust across sites, and replicates showed the expected increase in deamination with a 10-fold change in enzyme:substrate ratio (**Supplementary Figure 3c-d**). Given our interest in determinants of preferred sequences within DNA or RNA, we initially selected enzyme:substrate ratios that showed partial overall deamination. For ssDNA substrates reacted with 0.06 nM A3A, average editing across all cytosine bases was 12% (**Figure 3c, Supplementary Table**). Deamination could be readily detected across most other sites, with 89% of cytosines deaminated above background levels (>0.7%). The most deaminated site in ssDNA substrates, with 51% editing, was a cytosine at position 236, located in a predicted 3-nucleotide stem-loop within a TpC motif (**Figure 3d, Supplementary Figure 2b**). This correlates with a selectivity factor, defined as deamination at the most preferred site over average editing of sites throughout the substrate, of 4.3 (51% at site 236/12% overall all sites). Moreover, we observed that 100% of Cs in a 5’-T<u>C</u> context and 100% of Cs in a 5’-C<u>C</u> context were deaminated above background, along with detectable levels of deamination in the majority of 5’-R<u>C</u> sequence contexts (5’-A<u>C</u>, 71%; 5’-G<u>C</u>, 79%) (**Figure 3e-f**). While many of the highly targeted sites were predicted to occur in the loop of a stem-loop, we also noted several sites where the target cytosine is at the most proximal part of a stem adjacent to a loop on the 3’-side (**Supplementary Figure 2c**). This result suggests that dynamics at the end of a stem can impact A3A accessibility, a feature that has not been specifically detected before.

We then analogously examined the deamination of RNA reacted with 60 nM A3A, which is 1000-fold more enzyme than initially characterized with the ssDNA substrate. We observed that the average editing at sites in RNA was 2.6% (**Figure 3c**). Thus, while the restriction analysis at a known RNA hotspot (SDHB) suggested that RNA deamination was ~100-fold less efficient than DNA, when integrating over all sites, deamination was ~4000-fold less efficient on the matched RNA and DNA sequences. Interestingly, the cytosine that was most highly targeted in the RNA substrate differed from that of DNA. Rather than position 236, the preferred target in RNA was position 161, the SDHB target site, with 67% deamination and a selectivity factor of 26 (**Figure 3d**). The change in preferred target in DNA versus RNA is particularly intriguing in the context of prior work looking at the genome- and transcriptome-wide preferences suggesting that a 3-nucleotide loop is optimal for DNA substrates, while a 4-nucleotide loop (as in the SDHB site) is optimal for RNA substrates.^22–24^ Our results demonstrate that these alternative preferences, initially suggested by cell-based analysis, can be also detected *in vitro* with matched ssDNA and RNA substrates, and therefore likely reflect the intrinsic selectivity of A3A.

In striking contrast to the DNA substrate, in which deamination could be detected at most sites, only 50% of cytosines were deaminated above background levels (>0.7%) in RNA, suggesting that RNA deamination is more specific (**Figure 3c)**. Examining the sequence context preferences offers further support for altered selectivity in RNA versus DNA. We detected deamination above background in the majority of Cs in a 5’-T<u>C</u> or 5’-C<u>C</u> contexts, with deamination nearly undetectable in 5’-R<u>C</u> sequence contexts. To determine whether the increased impact of sequence context on deamination efficiency was consistent across different enzyme:substrate conditions, we repeated the amplicon sequencing analysis using varying A3A concentrations. Using these conditions, the difference between optimal and less-optimal substrates was greater for RNA than for DNA (**Figure 3f**, **Supplementary Figure 3e-f**). The existence of few highly edited sites in RNA and many more moderately deaminated sites in DNA supports the conclusion that RNA deamination occurs with low proficiency, but higher selectivity.

Taken together, our results indicate that A3A activity on ssDNA is generally efficient and broad, while its activity on RNA is greatly reduced and yet more selective across the 720-mer. Our analysis of the chimeric S35-rC substrate indicates that the target base itself drives a part of the nucleic acid selectivity (110-fold), but the rest of the RNA backbone also contributes considerably, explaining the overall ~4000-fold discrimination against RNA. Notably, RNA deamination requires ideal features – sequence context and secondary structure – that reflect selectivity, while DNA deamination shows a preference for similar features, but to a much lower extent. These observations also carry practical implications for efforts focused on understanding the activity of A3A in cancer mutagenesis or off-target base-editing activity. Specifically, rather than requiring transcriptome-wide analysis, our studies support the concept of focused analysis on highly edited RNA substrates such as the SDHB target as a more efficient approach to profiling A3A activity.^24^ Furthermore, given A3A’s patterns of selectivity for DNA versus RNA, off-target DNA activity and off-target RNA activity requires looking at distinct target genes or transcripts.

### Engineering A3A perturbs the balance of ssDNA and RNA editing

The use of DNA deaminases in concert with CRISPR/Cas proteins for targeted base editing has placed new emphasis on nucleic acid selectivity. Recent efforts have focused on mutating the deaminase to reduce undesirable RNA off-target mutagenesis.^14,30^ A base editor containing A3A with an N57G mutation (eA3A) was initially selected to increase the precision of cytosine editing in the targeting window but was also observed to significantly limit off-target RNA deamination.^14,15^ While eA3A appears promising in a base editor context, no direct biochemical study of the isolated eA3A domain acting on DNA or RNA has been reported. Given our observations of selectivity with the wild-type (WT) A3A, we hypothesized that enzymatic alterations that shift selectivity for DNA versus RNA might come with associated tradeoffs in specificity. As such, we sought to investigate substrate discrimination by eA3A using our matched 720-mer ssDNA and RNA assay (**Figure 3a**).

As with our examination of WT A3A, we first investigated deamination events at the SDHB hotspot via restriction enzyme digestions of amplicons generated from RNA and ssDNA substrates upon reaction with eA3A. Digestion of ssDNA-derived amplicons with ClaI showed only a ~5-fold increase in EC_50_ relative to WT A3A indicating that ssDNA editing was robust at the SDHB site (**Figure 4a**). By contrast, RNA editing was detectable, but only reaching 32% at the highest concentration evaluated (15 μM), suggesting a more dramatic reduction in RNA editing relative to the reduction observed with WT A3A. A similar pattern was observed when analyzing the DraI cleavage site. The consecutive deamination at this site could only be detected with the DNA substrate and the EC_50_ increased 55-times relative to WT A3A (**Supplementary Figure 4a**).

**Figure 4.**
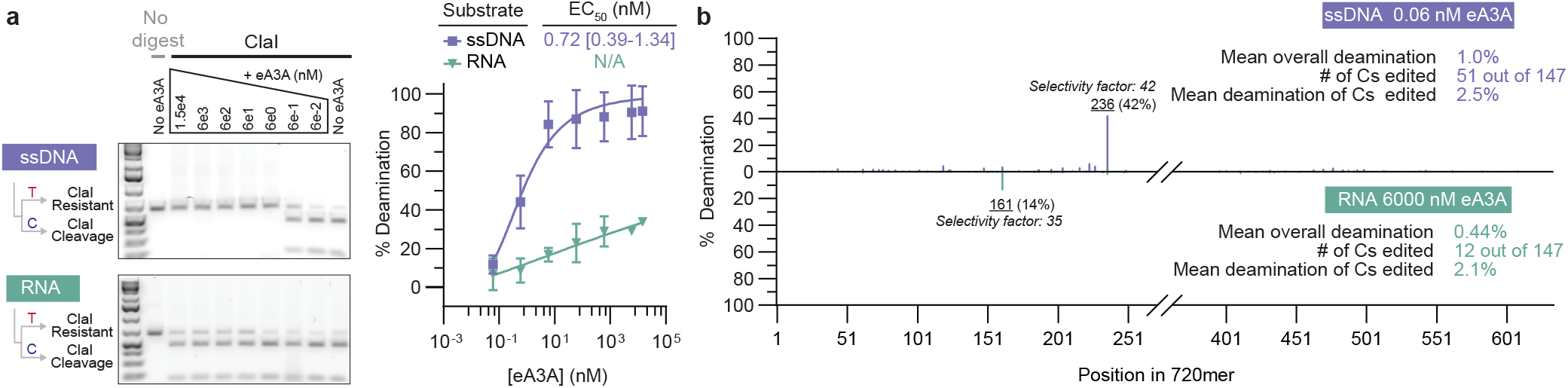
eA3A activity on long, single-stranded substrates with matched sequences. (a) Left – 10 ng of ssDNA or RNA substrates were reacted with eA3A (left to right, 15 μM and then 10-fold dilutions from 6 μM to 60 pM) for 30 min at 37 °C. Following PCR or RT-PCR, the amplicons were digested with ClaI, with representative gel images shown. Right – Quantification of percent deamination as a function of A3A concentration for ssDNA (purple) or RNA (green). Data represent mean and standard deviation from four independent replicates. (b) Base resolution map showing percent deamination vs. the position of cytosines across the 720mer as per NGS analysis. ssDNA data from reaction with 0.06 nM eA3A is shown in purple above the axis, while data from RNA reacted with 6 μM eA3A is shown in green below the axis. Data represent the mean deamination at each position from two independent experiments with results from individual amplicons provided in **Supplementary Table**, and the most heavily deaminated cytosine for each substrate labelled with its position, selectivity factor, and percent editing.

Moving from site-specific analyses to the broader amplicon, we next analyzed eA3A deamination by NGS. While the most targeted ssDNA site was still the stem-loop at position 236, we detected an overall pattern that appeared distinct from that observed with WT A3A. The overall ssDNA deamination was only 1.0% at 0.06 nM eA3A (**Figure 4b**), and 8.5% at 0.6 nM eA3A (**Supplementary Figure 4b**). Given the observed 12% deamination of ssDNA with 0.06 nM WT A3A, this result suggests that ssDNA deamination is decreased ~12-fold overall. Strikingly, however, editing at the position 236 hotspot is not reduced proportionally to others, showing deamination at 42% and yielding a selectivity factor of 42, a marked increase relative to the selectivity factor of 4.5 observed with WT A3A. Under higher enzyme:substrate ratios, the overall deamination across sites increases, but the preferred status for the 236 position remains notable and stands in contrast to WT A3A. Thus, the mutation introduced in eA3A has a small impact on activity at preferred sites but makes the enzyme far more selective when acting on ssDNA.

We next characterized RNA deamination across the amplicon using 6 μM eA3A. Here, we observed 14% editing at position 161, the SDHB hotspot, which was the only site deaminated >2.1% (**Figure 4b**). Notably, the change in hotspot targeting from position 236 in ssDNA to position 161 in RNA was consistent between WT A3A and eA3A, highlighting that selectivity factors differ between nucleic acid targets. However, while we observed 67% deamination of the SDHB site with 60 nM WT A3A, the 14% deamination observed with 6 μM eA3A leads us to estimate that, even at this preferred hotspot, RNA deamination is ~500-fold slower with eA3A relative to A3A. Since the preferred ssDNA hotspot at position 236 was deaminated nearly as efficiently by eA3A as with WT A3A, we conclude that the N57G mutation in eA3A enhances enzyme selectivity for both the nucleic acid target (DNA over RNA) and for mesoscale features that dictate preferred substrates.

Our results suggest that targeted active site manipulation with eA3A leads to modest decreased overall DNA reactivity (~12-fold), and that this altered activity manifests with a narrowed substrate scope for ssDNA (increased selectivity) and an even further narrowing of substrate tolerance for RNA. Remarkably, in the context of base editors, eA3A has been shown to target genomic loci with an efficiency that rivals that of WT A3A-containing base editors.^14^ It is likely that tethering of the DNA deaminase near the genomic target generated by Cas9 binding may permit even disfavored DNA targets to be deaminated effectively, while minimizing off-target activity on RNA.

## CONCLUSIONS

The purposeful deamination of DNA by A3A serves roles in retroviral restriction and in targeted genome editing. However, the nature of ssDNA engagement by the enzyme offers RNA as a possible alternative target that can result in unwanted pathological mutations. In this study, we have probed the mechanistic basis for A3A’s nucleic acid selectivity. From the nucleic acid angle, we show that the target cytidine plays a key role in selectivity, as the addition of a single 2’-OH at the target nucleotide’s sugar leads to a ~100-fold discrimination against RNA. Zooming out to both sequence context and secondary structure – which have been shown to play roles in DNA targeting – we demonstrate that these mesoscale features are more critical for RNA selectivity. RNA deamination is observed almost exclusively at sites with preferred sequence context and secondary structure, while DNA deamination can be observed when these features are non-ideal as well. The tradeoff between activity and specificity extends from nucleic acid determinants to those of the enzyme, as an active site mutation that lowers global deamination activity led to a disproportionate loss of RNA reactivity. Our conclusion that the preferred target in DNA can differ from that in RNA offers important insights for the design of optimal reporter substrates for tracking A3A’s specific activity on DNA or RNA, or its global activity in cells. These insights also highlight the fact that optimization of A3A activity to enhance DNA selectivity or even engineer RNA selectivity may be possible,^14,31^ but would be anticipated to come at a cost in the breadth of activity.

## MATERIALS AND METHODS

### A3A and eA3A expression

The A3A expression construct (Addgene #109231) has been previously described and can be used to purify A3A as a fusion protein (MBP-A3A-His) that can be further processed to generate the isolated A3A domain.^32,33^ For eA3A (A3A-N57G), the N57G mutation was introduced via Q5 site directed mutagenesis (New England Biolabs, NEB). Bacterial expression of A3A and eA3A constructs has been previously described in detail.^33^ Purified MBP-A3A-His, MBP-eA3A-His or isolated A3A were dialyzed overnight in 50 mM Tris-Cl (pH 7.5), 50 mM NaCl, 10% glycerol, 0.5 mM DTT and 0.01% Tween-20 and the concentrations of proteins were determined using a BSA standard curve.

### SwaI-based deaminase activity on ssDNA and chimeric substrates

5’-fluorescein (FAM) fluorescently labelled substrate S35-dC or a matched substrate with a single target ribocytosine in an otherwise DNA backbone (S35-rC) were synthesized by Integrated DNA Technologies (IDT), along with the associated product controls (S35-dU and S35-rU). 100 μM oligonucleotide was treated with 6-fold dilutions of untagged-A3A (from 1 μM to 4 pM) in optimal A3A reaction conditions (final, 20 mM succinic acid:NaH_2_PO_4_:glycine (SPG) buffer pH 5.5, 0.1% Tween-20). The reaction was allowed to proceed for 30 min at 37 °C and then terminated (95 °C, 10 min). 200 nM of the complementary strand was then added and annealed. SwaI (NEB) was added, and digestion carried out overnight at room temperature. Formamide loading buffer was added and samples were heat denatured (95 °C, 20 min), and then run on a 20% denaturing TBE/urea polyacrylamide gel at 50 °C. Gels were imaged using FAM filters on a Typhoon imager (GE Healthcare). Area quantification tool in ImageJ was used for quantitative analysis.

### Synthesis of 720-mer ssDNA substrates

To generate ssDNA, a 720-bp gBlock gene fragment (IDT) was used as a template (**Supplementary Figure 2a**) and amplified with Taq polymerase (NEB) using a linear-after-the-exponential(LATE)-PCR reaction protocol, which employs excess forward primer relative to a phosphorylated reverse primer.^32^ The reactions were purified (NucleoSpin, Fisher) and then treated with λ exonuclease for 1 h at 37 °C to degrade the phosphorylated strand, followed by heat inactivation (90 °C, 10 min). The products were then run on a 2% agarose gel and the ssDNA was recovered by using Gel DNA Recovery Kit (Zymoclean). The ssDNA was further purified by ethanol precipitation, and its concentration was measured using a Qubit^®^ fluorometer (ThermoFisher). For one replicate, ssDNA was obtained as a megamer oligonucleotide (IDT) and further purified by ethanol precipitation.

### Synthesis of 720-mer RNA substrates

Using the 720-bp gene block (IDT) dsDNA as a template, RNA was generated via *in vitro* transcription using TranscriptAid Enzyme Mix (ThermoFisher) under recommended conditions and incubated for two hours at 37 °C. The RNA was then purified via phenol-chloroform extraction and ethanol precipitation. The sample was resuspended in nuclease-free water and further treated with MspI, XbaI, and AclI restriction enzymes (NEB) to digest any remaining DNA template. After a 1 h incubation at 37 °C, the RNA Clean and Concentrator-5 kit (Zymo Research) was used to purify the RNA. To further ensure complete removal of template DNA, the RNA was treated with DNase I (Ambion) for 30 min at 37 °C. Purification was repeated (RNA Clean and Concentrator-5), and the concentration of purified RNA was measured using a Qubit^®^ fluorometer. Secondary structure of several mesoscale regions in the 720-mer are predicted via the “Predict a Secondary Structure Web Server” (https://rna.urmc.rochester.edu/RNAstructureWeb/Servers/Predict1/Predict1.html), and the structures with the lowest free energy prediction are provided in **Supplementary Figure 2c**.

### Qualitative deaminase activity of 720-mer using restriction enzyme-based method

10 ng of either the 720-mer ssDNA or RNA substrate were reacted with varied concentrations of MBP-A3A-His or MBP-eA3A-His in 20 mM SPG (pH 5.5) with 0.1% Tween-20 in a 10 μL total volume. 10U RNase inhibitor was added to the reaction mixtures of the RNA samples only. After 30 min at 37 °C, the reaction was terminated by denaturation at 95 °C for 10 min. RNA samples were reverse transcribed using M-MuLV Reverse Transcriptase (NEB) for 1 h at 42 °C, followed by heat inactivation (**Supplementary Figure 2b)**. 2 μL from both ssDNA and reverse-transcribed RNA samples were used to template PCR amplification using Taq polymerase (NEB). The resulting amplicons were treated with either ClaI or DraI (NEB) for 1 h at 37 °C for position-specific analysis. Samples were then run on either a 1% TAE or 1.5% TBE agarose gel, which were imaged on a Typhoon imager (GE Healthcare).

### Sequencing Data Analysis

Amplicons from the 720-mer assay were purified via the QIAquick PCR Purification kit (Qiagen). Concentrations were measured using a Qubit^®^ fluorometer (ThermoFisher) and then sequenced by Amplicon-EZ Next Generation Sequencing (Genewiz). Read qualities were evaluated by FastQC v0.11.9 (http://www.bioinformatics.babraham.ac.uk/projects/fastqc/). Low-quality sequence (Phred quality score <28) and adapters were trimmed via Trim Galore v0.6.5 (http://www.bioinformatics.babraham.ac.uk/projects/trim_galore/) prior to analysis with CRISPResso2.^34^ Sequencing analysis of amplicons generated from ssDNA processed in parallel but without any A3A were used to compute the background level of deamination in cytosine bases (estimated at 0.7%). The data shown represent the mean deamination at each position, averages from at two independent experiments. Results from individual amplicons are provided in **Supplementary Table**. Selectivity factor, a measure of deamination at a specific site relative to the average deamination across the 720mer, is calculated as shown below:

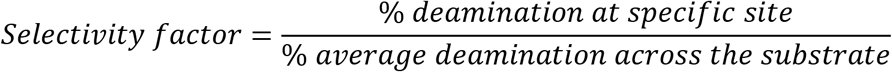

## ACKNOWLEDGMENTS

K.N.B. was supported by an individual fellowship from the National Science Foundation. This work was supported by the National Institutes of Health R01-GM138908 (to R.M.K.).

