## Supplementary Figures 1-4 for "The base-editing enzyme APOBEC3A catalyzes cytosine deamination in RNA with low proficiency and high selectivity"

* Co-first authors; † Corresponding author

**Supplementary Table.** Sequencing analysis of 720mer ssDNA and RNA substrates.

#### SUPPLEMENTARY FIGURE LEGENDS

**Supplementary Figure 1. Active site structure near sugar of dC.** (a) Co-crystal structure of A3A and ssDNA (PDB: 5SWW) depicting the U-shaped conformation of the substrate. (b) A zoomed in view of the target cytosine ($dC_0$) inserted in the active site. The residues implicated in $Zn^{2+}$ binding (H70, C101, C106) are shown as sticks. The N57 residue, altered in eA3A, is additionally shown.

**Supplementary Figure 2. 720mer substrate.** (a) Sequence of the 720mer DNA, including T7 promoter (annotated), primer set 1 (red) and primer set 2 (blue). (b) Secondary structure prediction of eight mesoscale regions (~30 bp) with relatively highly edited cytosines. The "Predict a Secondary Structure Web Server" was utilized (https://rna.urmc.rochester.edu/RNAstructureWeb/Servers/Predict1/Predict1.html), and the structure with the lowest free energy prediction is provided (except for site 119, where the second lowest free energy prediction is shown). (c) Following A3A reaction, RNA substrates were subjected to reverse transcription (RT). This is a representative gel depicting RNA substrates following the RT reaction (+) or without the RT reaction (-).

**Supplementary Figure 3. A3A activity on long, single-stranded substrates with matched sequences.** (a) 720mer assay diagram. Sequenced-matched ssDNA and RNA substrates were reacted with A3A. The samples were amplified by PCR (ssDNA) or RT-PCR (RNA). Amplified products were then subjected to site-specific examination via the use of either ClaI or DraI restriction enzymes. Notably, deamination renders the amplicon resistant to ClaI digestion and susceptible to DraI. (b) Left – Representative gels of A3A titration. 10 ng of ssDNA and RNA substrates were treated with 10-fold dilutions of A3A (6 µM to 6 pM from left to right) for 30 min at 37 ˚C. Following amplification, amplicons were digested with DraI and analyzed on a 1.5% agarose gel. Right – Quantification of percent deamination as a function of A3A concentration for ssDNA (purple) or RNA (teal). Data represent four independent replicates with mean and standard deviation plotted. Product formation was fit to determine the $EC_{50}$, the enzyme concentration required to convert half of the substrate to product under the assay conditions. (c) Correlation scatter plot showing percent deamination in ssDNA after treatment with 0.06 nM A3A vs. 0.6 nM A3A. The -1 position of the C to T converted sites are indicated by the different colors. A y = 0.1x line is shown for reference given the comparison of 10-fold A3A concentration difference in ssDNA activity. The data show the expected proportionate increase, whereby substrate with low deamination increase ~10-fold. (d) Correlation scatter plot showing percent deamination in RNA after treatment with 6 nM A3A vs. 60 nM A3A. The -1 position of the C to T converted sites are indicated by the different colors. A y = 0.1x line is shown for reference given the comparison of 10-fold A3A concentration difference in RNA activity. (e-f) Base resolution maps showing percent deamination vs. the position of cytosines across the 720mer as per NGS analysis for several conditions. The data represent the mean deamination at each position from two independent experiments with results from individual amplicons provided in **Supplementary Table**.

**Supplementary Figure 4. eA3A activity on long, single-stranded substrates with matched sequences.** (a) Left – 10 ng of ssDNA and RNA substrates were reacted with eA3A (left to right, 15 µM and then 10-fold dilutions from 6 µM to 60 pM) for 30 min at 37 ˚C. Following PCR or RT-PCR, the amplicons were digested with DraI, with representative gel images shown. Right –

Quantification of (a), showing percent deamination as a function of eA3A concentrations for ssDNA (purple) or RNA (green). Data represent mean and standard deviation from four independent replicates. Product formation for ssDNA was fit to determine the $EC_{50}$. (b-c) Base resolution maps showing percent deamination vs. the position of cytosines across the 720mer as per NGS analysis for several conditions. The data represent the mean deamination at each position from two independent experiments with results from individual amplicons provided in **Supplementary Table**.

#### Supplementary Figure 1

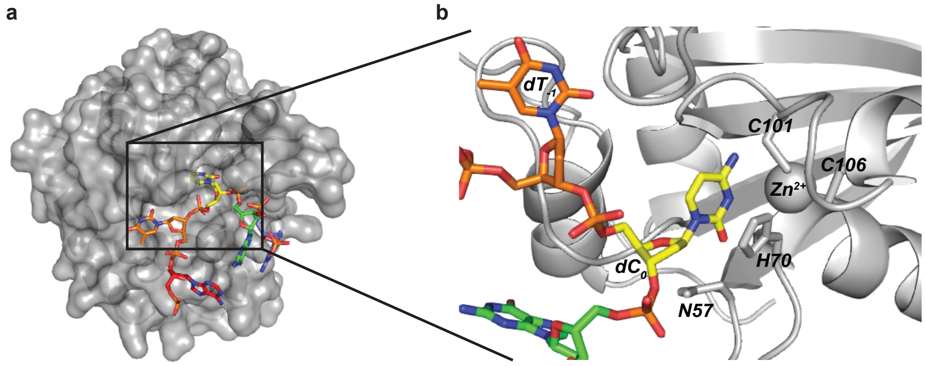

### Supplementary Figure 2

**a** 720mer DNA Sequence

T7 Promoter

#1

TAATACGACTCACTATAGGATGTATAGAATGATGAGTTAGGTAGTGTTGATATGGGTTATGAATGAAGTATCCGTTCATCAGACTCTATCA
GCCAGCCATGCGCCATCGTACACCGTTCATCGTCCGCTTCAAGTCAGTTCGATTCCCATGACCGTCTGCGCCTCGTTCCGGCTTAAC
AGAGCAGGTCGCATTCGACGCCATCTATCGATGGACTAATCAGGCATACATCTCCGTACTGTTCGCGCTATCGCTGGTCAATTCTTGC
CTCGTTCGGTGCCTTCGGCTTACTTACCCGAACTTCCACGGTCATCCTCAGCCTCTAGCCATTGCTACCCTCATTCCATGCCGTCATT
CGCAGCTTACGATCCCGCAAGCGGCCTAACGTCCCAGCAGCCTCAGCGACCGTTATCTTGCAGGCGATGTCATGTCGGCACAACTA
TCATATCATTGGATTCCAAATCCAGAGCGATTCACCTCAGCAGCTAACCGATACAAGCTCCTAGCCTTCAACGTTCGCATTCCATGTTC
CGTTCATTCTCACTCCGCTAACAGCAATCCCGCAACAGCATCTGTTGCATACATTCTACCGTCTAGACGACAACTTATCGTTACGCTAA
CTAGGGCTGTCATGCTACAGGCGTACTGACGAACTCAGTGTTAAGGTATATGAGTAGATGATTGATTGGGTATGTTGATAAGTGTA

#633

Primer set 1

Primer set 2

**b**

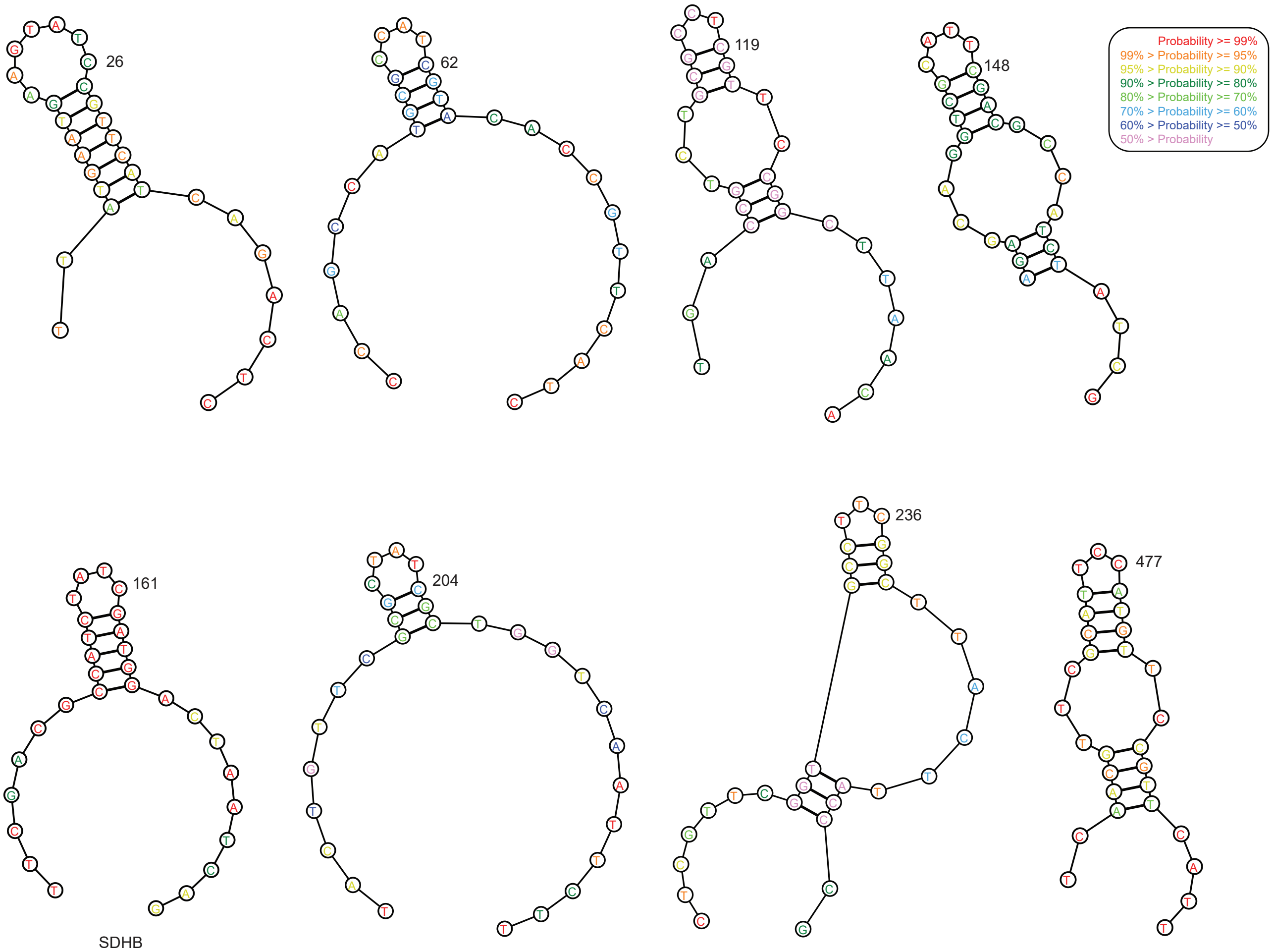

**c**

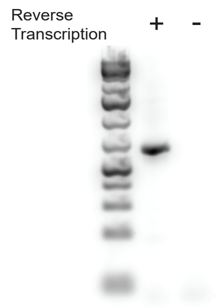

### Supplementary Figure 3

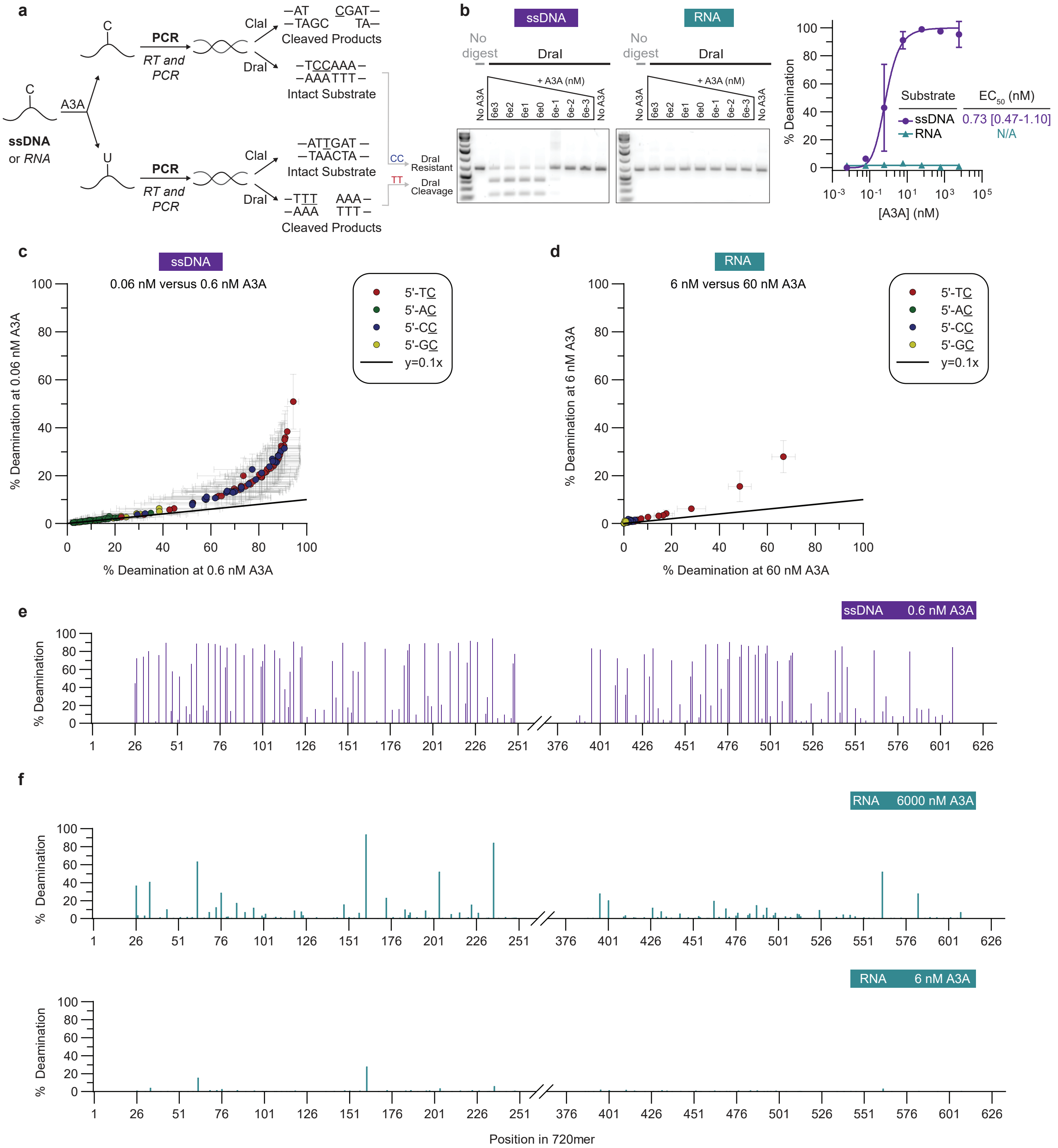

### Supplementary Figure 4

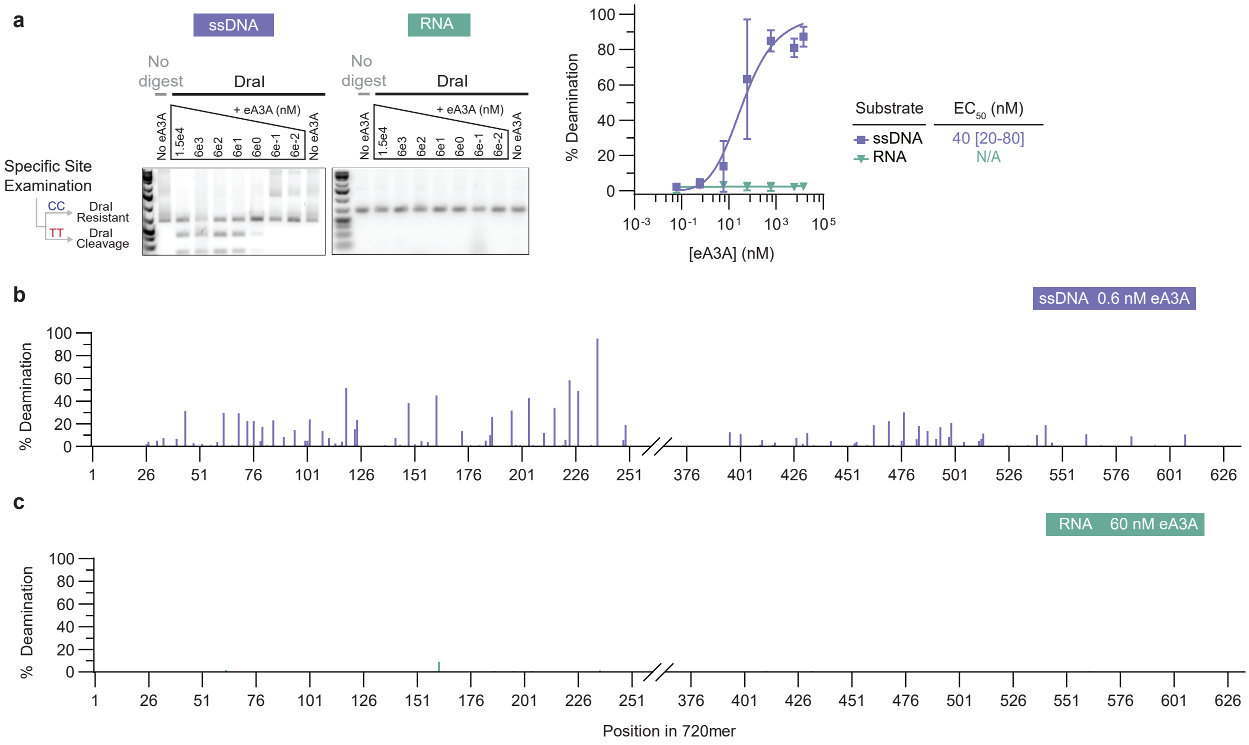
